# Matrix metalloproteinases proteolyze RAB proteins and contribute to cisplatin-induced ototoxicity

**DOI:** 10.64898/2026.02.28.708770

**Authors:** Zahra Zandi, Bridgette Hartley, Wesam Bassiouni, Michèle G. DuVal, Shu Luo, Maria J. Spavor, W. Ted Allison, Olivier Julien, Richard Schulz, Amit P. Bhavsar

## Abstract

Matrix metalloproteinases (MMPs) are rapidly expressed and activated in response to oxidative stress and contribute to various pathological conditions. Cisplatin is a highly effective chemotherapeutic agent; however, its clinical use is limited by its associated permanent hearing loss (ototoxicity). While cispwlatin-induced oxidative stress and inner ear cell death are well-established, the contribution of MMPs remains unclear. In this study, we demonstrate that cisplatin exposure triggers activation of MMP-2 and MMP-9 and expression of an intracellular N-terminal-truncated isoform of MMP-2 in mouse inner ear hair cells. Pharmacological inhibition of MMP-2 and genetic knockdown of *Mmp-9* enhanced hair cell survival and attenuated cisplatin-induced inflammation and cytotoxicity. Furthermore, proteomic analysis revealed that proteins involved in intracellular trafficking, including RAB proteins, may serve as potential substrates of intracellular MMP-2 upon cisplatin exposure, pointing to a previously unrecognized mechanism of cisplatin-induced hair cell injury. *In vitro* analysis confirmed that MMP-2 cleaves RAB9A in response to cisplatin, and *in silico* analyses predicted MMP-2-preferred cleavage sites on RAB9A. Collectively, our findings identify MMP-2 as a promising therapeutic target for mitigating cisplatin-induced ototoxicity.

## Introduction

Cisplatin is a widely used chemotherapeutic agent for the treatment of a broad spectrum of adult and pediatric solid tumors, including head and neck, ovarian, and lung cancers in adults, as well as neuroblastoma, osteosarcoma, germ cell tumors, hepatoblastoma, brain tumors, and retinoblastoma in children (Dasari and Tchounwou 2014; Ruggiero et al. 2013). Cisplatin has markedly improved 5-year overall survival rates to as high as 80%, making it an indispensable component of the treatment regimen for many types of cancer (Chua et al. 2005; Kleinberg et al. 2003; Stoter et al. 1984). The anti-tumor activity of cisplatin stems from its ability to form guanine cross-links in DNA, leading to cell cycle arrest and cell death (Cohen and Lippard 2001; Dasari and Tchounwou 2014). However, cisplatin also accumulates in non-tumor tissues such as the kidneys, neurons, and inner ear, causing dose-limiting toxicities including nephrotoxicity, neurotoxicity, and ototoxicity, i.e., hearing loss (Qi et al. 2019). Among these, cisplatin-induced ototoxicity (CIO) is particularly debilitating. CIO manifests as permanent, progressive, bilateral high-frequency hearing loss affecting 40-80% of patients, with children being especially vulnerable (Dai et al. 2024). At higher doses (150–225 mg/m²), CIO can occur in nearly all patients, limiting the maximal tolerable dose of cisplatin and ultimately compromising its anti-cancer efficacy (Kopelman et al. 1988; Chang and Chinosornvatana 2010). In children, the consequences of CIO are especially severe, leading to long-term impairments in neurocognitive development, speech and language acquisition, and psychosocial well-being (Knight et al. 2005).

Cisplatin preferentially and indefinitely accumulates in key auditory structures of the cochlea of the inner ear, including the organ of Corti, stria vascularis, and spiral ganglion neurons, leading to progressive cell death (Rybak 2007; Breglio et al. 2017). The mechanosensory outer hair cells in the organ of Corti, which are responsible for sound transduction to the central nervous system, are of particular concern regarding cisplatin-induced damage since these cells cannot regenerate after injury (Lim 1986). Among the best-characterized mechanisms underlying CIO is cell death driven by the overproduction of reactive oxygen and nitrogen species (RONS) in the cochlea (Clerici and Yang 1996; Lee et al. 2004). Sodium thiosulfate, a RONS scavenger, was approved by the FDA in 2022 for CIO prevention; however, its use is limited to non-metastatic cases, and its protective efficacy is suboptimal (Freyer et al. 2017). These challenges highlight the urgent need to develop more effective and broadly applicable strategies to prevent or mitigate CIO.

Matrix metalloproteinases (MMPs) are a family of zinc-dependent proteolytic enzymes that regulate various aspects of physiological cellular processes and tissue functions, including cell differentiation, proliferation, angiogenesis, and wound healing, especially by cleaving extracellular matrix proteins (Bassiouni et al. 2021; Cabral-Pacheco et al. 2020). Conversely, the dysregulation of these proteases has been implicated in numerous pathological conditions, including cardiovascular disorders and cancer (Bassiouni et al. 2021). Importantly, intracellular isoforms of MMPs also cleave various intracellular substrates during pathological conditions, such as MMP-2 during myocardial ischemia-reperfusion injury (Wang et al. 2002). The activity of intracellular MMPs, particularly MMP-2 and MMP-9, has been implicated in various subcellular compartments, such as mitochondria and nuclei (Hughes et al. 2014; Lee et al. 2021). MMPs consist of an N-terminal prodomain and a catalytic domain. In the inactive state, a cysteine residue within the prodomain coordinates with the zinc ion in the catalytic site, preventing substrate binding. Full enzymatic activation requires the removal of this inhibitory prodomain through either proteolytic cleavage or non-proteolytic mechanisms (Nagase et al. 2006). Oxidative stress can rapidly activate full-length MMPs via a non-proteolytic mechanism, in which oxidation of a cysteinyl thiol group in the prodomain triggers a conformational change that exposes the catalytic site and permits substrate binding (Okamoto et al. 2001; Viappiani et al. 2009).

In addition, oxidative stress induces the expression of a unique intracellular MMP-2 isoform lacking the first 76 amino acids, termed N-terminal truncated MMP-2 (NTT-MMP-2). This transcriptional variant lacks both the signal peptide and the prodomain, resulting in intracellular retention and constitutive activity. NTT-MMP-2 expression is strongly induced by oxidative stress and plays an important role in regulating innate immune responses (Lovett et al. 2012). Moreover, full-length MMP-2 can also be retained within cells due to the inefficiency of its signal peptide, where it becomes activated and degrades intracellular targets (Ali et al. 2012).

Despite the well-established contribution of RONS to cisplatin-induced ototoxicity (Yin et al. 2017; Kim et al. 2009), the specific roles of full-length and truncated MMP-2 and MMP-9 in this process have not been investigated. In this study, we report for the first time that cisplatin exposure upregulates both the expression and enzymatic activity of MMP-2 and MMP-9 in the mouse outer hair cell line HEI-OC1. Our data further demonstrate that these MMPs contribute to inflammatory signaling and cell death. Importantly, pharmacological inhibition of MMP-2 significantly reduced cisplatin-induced cytotoxicity, underscoring its potential as a therapeutic target for preventing or mitigating CIO. Proteomic analyses revealed several potential MMP-2 substrates, including proteins involved in intracellular transport, such as Ras-associated binding proteins (RABs), suggesting a previously unrecognized mechanism by which MMP-2 may contribute to cochlear pathology.

## Results

### MMP-2 and MMP-9 are activated in response to cisplatin

Since MMPs are activated by oxidative stress, and RONS play a key role in cisplatin-induced toxicity (Santos et al. 2007; Kim et al. 2009; Yin et al. 2017), we aimed to investigate whether cisplatin activates extracellular or intracellular MMP-2 and MMP-9 in mouse auditory hair cells (HEI-OC1). MMP-2 activity was measured in the conditioned media of HEI-OC1 cells after 6 hours of treatment with 20 μM cisplatin using gelatin zymography. As shown in Fig. 1A and B, 6 hours of cisplatin treatment resulted in a small but significant increase in the intensity of the 72 kDa band, corresponding to MMP-2, compared to the untreated control. In contrast, MMP-9 activity was not detected in the conditioned media of either group at this time point. However, MMP-9 activity became detectable after 24 and 48 hours of cisplatin treatment (Supplementary Fig. 1A, B) and (Fig. 1C, D), with a significant, concentration-dependent increase in the 92 kDa band observed at the 48-hour time point (Fig. 1C, D).

**Figure 1.**
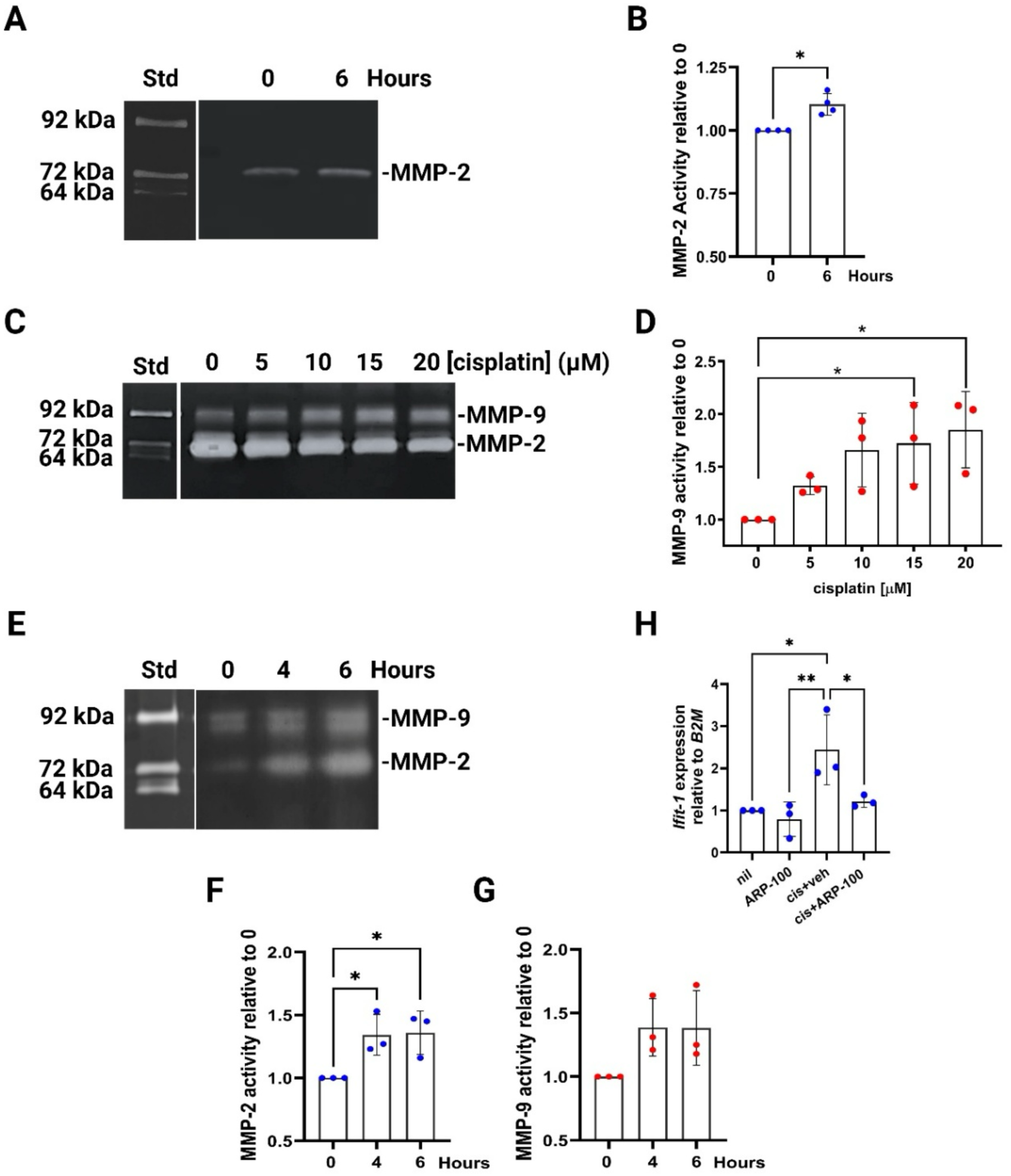
MMP-2 and MMP-9 are activated in response to cisplatin. Gelatin zymography (A, C, E) and corresponding quantitative analysis (B, D, F, G) of MMP-2 and MMP-9 activities in the conditioned media of HEI-OC1 cells following 6 h (A, B) and 48 h (C, D) of exposure to 20 μM cisplatin. Cytosolic MMP-2 and MMP-9 activities were measured following 4 h and 6 h of treatment with 20 μM cisplatin (E-G). Standard (Std) derives from conditioned medium from HT-1080 cells. H) mRNA expression of *Ifit-1* was measured using RT-qPCR following a 24h exposure to 20 μM cisplatin in the presence or absence of ARP-100 (5 μM), compared to the untreated control (nil), and normalized to *B2M* expression. The data are represented as the mean ± SD of three independent experiments. Statistical significance was determined using one-way ANOVA followed by Dunnett’s post-hoc test (\**p* < 0.05 and \*\**p*<0.01).

To investigate intracellular MMP activity, mitochondrial and cytosolic fractions of HEI-OC1 cells were isolated following cisplatin exposure. As shown in Fig. 1E-G, intracellular MMP-2 activity was significantly elevated after 4 and 6 hours of treatment with 20 μM cisplatin. However, the increase in intracellular MMP-9 activity was not statistically significant. Notably, no increase in the activity of either MMP was observed in the mitochondrial fraction (data not shown). The N-terminal truncated (NTT) MMP-2, an intracellular isoform of MMP-2, is upregulated in response to oxidative stress and plays a role in modulating innate immune responses (Lovett et al. 2012). It has been shown that NTT-MMP-2 specifically induces the transcriptional upregulation of IFIT-1, a regulator of ribosomal translation (Lovett et al. 2012). Interestingly, the result of real-time quantitative PCR (RT-qPCR) showed that cisplatin significantly upregulated *Ifit-1* gene expression, an effect that was reversed by the MMP-2 inhibitor, ARP-100 (Fig. 1H), highlighting that cisplatin induced the expression of *NTT-Mmp-2* and its downstream mediators.

Moreover, RT-qPCR indicated that the expression of *NTT-Mmp-2* increased ∼6-fold within 1 hour of treatment with 20 μM cisplatin before declining at later time points (Fig. 2A). At the same time, no significant changes were detected in full-length (FL) *Mmp-2* expression (Fig. 2B). In contrast, *Mmp-9* expression showed a ∼3-fold increase after 2 hours of exposure (Fig. 2C). Immunofluorescence staining confirmed the presence of intracellular MMP-2 and MMP-9 throughout the cells and showed an increase in fluorescence signal following 6 h of cisplatin treatment, especially inside and around the nucleus (Fig. 2D and Supplementary Fig. S1C). Of note, the MMP-2 activity observed in the cytoplasm could originate from either the nascent constitutively active NTT isoform produced in response to oxidative stress or the full-length isoform that has been activated. Taken together, these results indicate that both full-length and truncated MMP-2 are rapidly activated following cisplatin exposure, whereas secreted MMP-9 is activated later.

**Figure 2.**
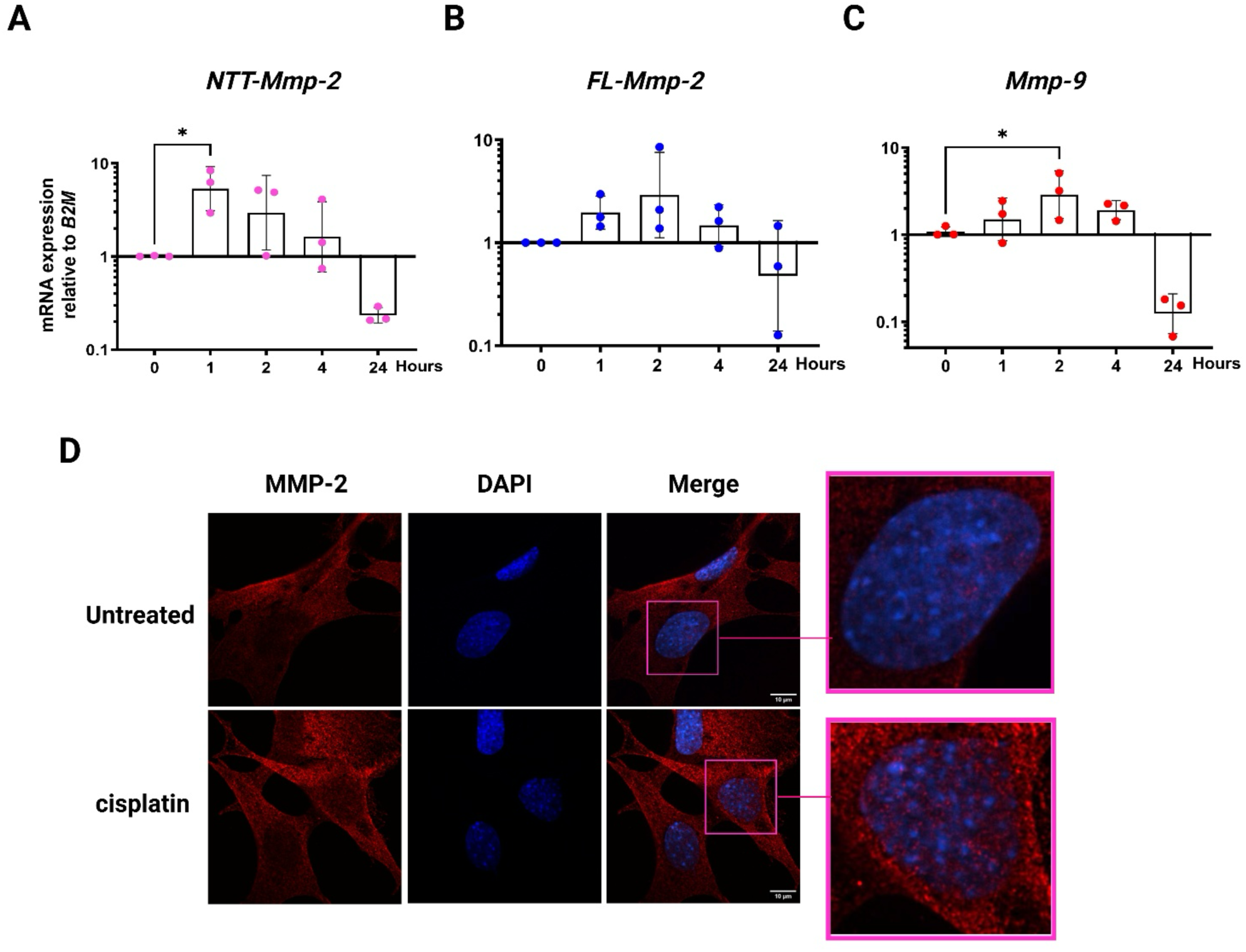
Enhanced *Mmp-2* and *Mmp-9* expression in response to cisplatin treatment in HEI-OC1 cells. mRNA expression of N-terminal truncated (NTT)-*Mmp-2* (A), full-length (FL)-*Mmp-2* (B), and *Mmp-9* (C) was measured using RT-qPCR following exposure to 20 μM cisplatin for various timepoints, compared to the untreated control (0), and normalized to *B2M* expression. D) Immunofluorescence staining of HEI-OC1 cells treated with 20 μM cisplatin for 6 h or untreated cells. Staining was performed using an antibody specific to MMP-2, followed by detection with a secondary antibody conjugated to Alexa Fluor 647 (red). The zoomed-in boxes indicate nuclei with an increased red signal (magenta). The data are represented as the mean ± SD of three independent experiments. Statistical significance was determined using one-way ANOVA followed by Dunnett’s post-hoc test (\**p* < 0.05).

### MMP-2 preferring inhibitors attenuate cisplatin toxicity

Given the observed upregulation and activation of MMP-2 following cisplatin treatment, we sought to determine its contribution to cisplatin-induced toxicity in HEI-OC1 cells. To this end, we performed a cisplatin concentration response using an MTT assay as a proxy for cell viability in the presence of vehicle control or one of two orally active MMP-2-preferring inhibitors, ARP-100 and ONO-4817 (Yamada et al. 2000; Rossello et al. 2004). We observed that the IC_50_ of cisplatin increased from 33 to 73 μM when co-treated with 10 μM ARP-100 (*p* = 0.0002). Similarly, co-treatment of cisplatin with 10 μM of ONO-4817 increased the IC_50_ from 33 to 126 μM (*p* = 0.0002) (Fig. 3A, B). These experiments indicate that MMP-2 inhibition significantly mitigates cisplatin-induced cytotoxicity.

**Figure 3.**
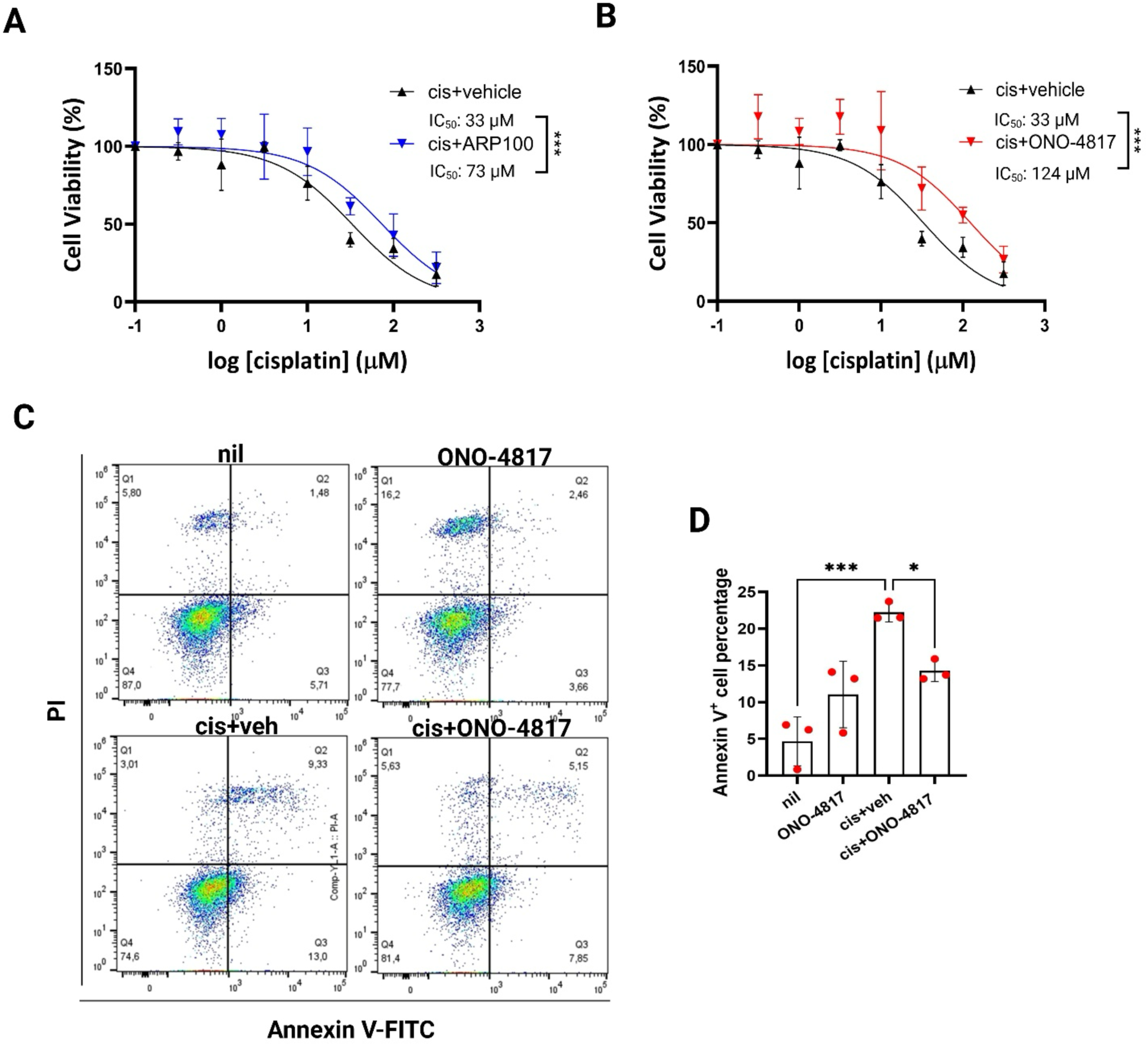
MMP-2-preferring inhibitors attenuate cisplatin toxicity in HEI-OC1 cells. Cell viability of HEI-OC1 was assessed by MTT assay following a 24 hour treatment with cisplatin in the absence or presence of 10 μM ARP-100 (A) or ONO-4817 (B). (C,D) Representative flow cytometry (C) and quantitative analysis (D) of AnnexinV^+^-cells (Q2 & 3) following a 24 hour cisplatin (20 μM) exposure in the absence or presence of ONO-4817 (5 μM) or vehicle; nil represents the untreated control. The data are represented as the mean ± SD of three independent experiments. Statistical significance was determined using one-way ANOVA followed by Dunnett’s post-hoc test (\**p* < 0.05 and \*\*\**p* < 0.001). Statistical significance in A and B was calculated from the comparison between IC_50_ values of the inhibitors and the vehicle.

To further assess the effect of MMP-2 inhibition on cisplatin-induced apoptosis, flow cytometry was performed to quantify Annexin V and PI markers in cells treated with cisplatin and the MMP-2 inhibitors (Fig. 3C, D). This analysis revealed that treatment with 20 μM cisplatin induced approximately 22% apoptotic cells after 24 hours of treatment (Annexin V^+^-cells). Cells pre-treated with 5 μM ONO-4817 followed by treatment with the same concentration of cisplatin showed a 56% decrease in the population of Annexin V^+^/PI^-^ cells, suggesting a protective effect. Notably, ONO-4817 treatment alone led to a marginal increase in apoptosis, consistent with studies reporting the physiological role of MMP-2 in cochlear cell survival under normal conditions (Wu et al. 2017; Sung et al. 2014).

### MMP-2 modulates cisplatin-induced interleukin-6 secretion

The secretion of the pro-inflammatory cytokine IL-6 is a well-established indicator of cisplatin-induced toxicity in cochlear cells, as it promotes the generation of RONS and triggers cell death (Babolmorad et al. 2021; Ramkumar et al. 2021; Tan and Vlajkovic 2023). While the roles of MMP-2 and MMP-9 in modulating inflammatory cytokines and regulating innate immune responses are well-documented (Song et al. 2013; Lovett et al. 2012), their direct involvement in inflammation within cochlear cells remains unclear. To investigate the potential role of MMP-2 in cisplatin-induced IL-6 secretion, HEI-OC1 cells were transfected with either an MMP-2-expressing plasmid or an empty vector (EV), followed by treatment with cisplatin or PBS. The overexpression and activation of MMP-2 were confirmed in this cell model (Supplementary Fig. 2A, B). As shown in Fig. 4A, cisplatin alone induced a ∼3-fold increase in IL-6 levels, while MMP-2 overexpression alone resulted in a ∼4-fold increase compared to EV-transfected, PBS-treated controls. Notably, cells overexpressing MMP-2 and treated with cisplatin exhibited a significantly greater induction of IL-6 than either MMP-2-overexpressing cells treated with PBS or EV-transfected cells treated with cisplatin. These findings suggest an additive interaction between MMP-2 and cisplatin in promoting IL-6 secretion, supporting a role for MMP-2 in amplifying cisplatin-induced inflammatory responses in cochlear cells.

**Figure 4.**
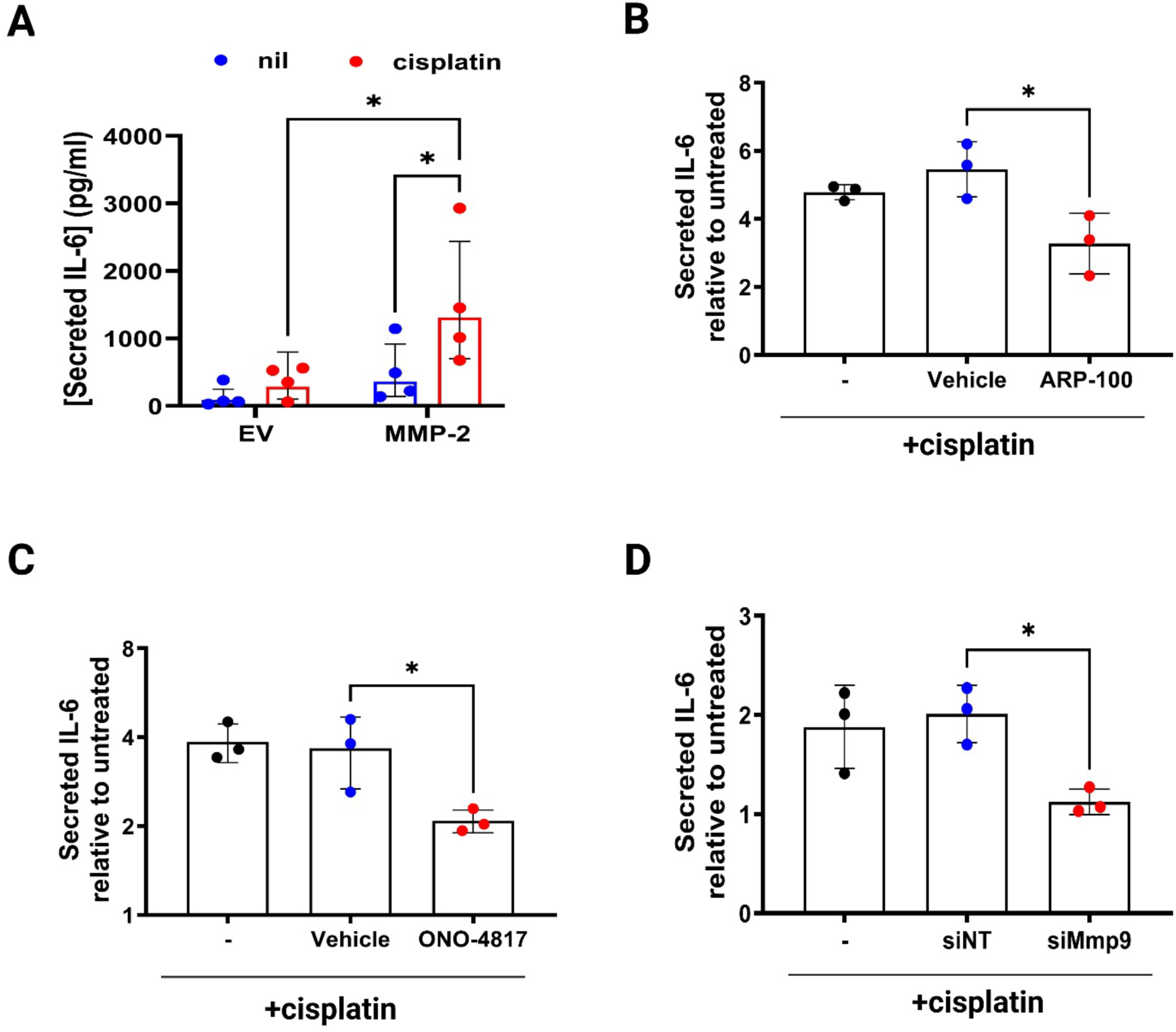
MMP-2 modulates cisplatin-induced interleukin-6 secretion in HEI-OC1 cells. (A) IL-6 secretion in cells transfected with either an empty vector (EV) or an *Mmp-2*-expressing vector and treated with 100 μM cisplatin or left untreated (nil) for 24 h. (B, C) IL-6 secretion in cells pre-treated for 2 h with the vehicle or 5 μM ARP-100 (B) or ONO-4817 (C), followed by treatment with 20 μM cisplatin for 24 h. (D) IL-6 secretion in cells transfected with non-targeting siRNA (*siNT*), *Mmp-9*-targeting siRNA (*siMmp-9*), and exposed to 20 μM cisplatin for 24 h. The data are represented as the mean ± SD of three independent experiments. Statistical significance was determined using one-way ANOVA followed by Dunnett’s post-hoc test (\**p* < 0.05).

Conversely, MMP-2 preferring inhibitors attenuated cisplatin-induced IL-6 secretion. As shown in Fig. 4B, C, co-treatment with 5 μM of either ARP-100 or ONO-4817 significantly reduced IL-6 levels by half compared to cisplatin combined with vehicle, further highlighting the contribution of MMP-2 to the inflammatory response. Moreover, the mRNA expression of *Il-6* was induced by 24-hour cisplatin treatment, while co-treatment with ARP-100 significantly reduced *Il-6* expression (Supplementary Fig. 3C).

Due to the lack of commercially available selective small-molecule inhibitors for MMP-9, gene silencing was used to evaluate its role in CIO. HEI-OC1 cells were transfected with siRNA targeting *Mmp-9* (*siMmp-9*) or a non-targeting control (*siNT*) prior to cisplatin treatment. As shown in Fig. 4D, *siMmp-9*-transfected cells exhibited nearly a 50% reduction in IL-6 secretion compared to the *NT-siRNA*-transfected group, indicating that *Mmp-9* expression also plays a significant role in cisplatin-induced inflammation in HEI-OC1 cells.

### MMP-2 preferring inhibitors do not interfere with the anti-cancer activity of cisplatin

An effective otoprotective therapy should protect auditory hair cells from cisplatin-induced damage without substantially compromising its antitumor efficacy. To assess whether MMP-2 inhibition interferes with the cytotoxic activity of cisplatin, we evaluated the combined effects of cisplatin and MMP-2 inhibitors on cancer cell viability *in vitro*. Cisplatin concentration-response was measured using an MTT assay in the presence of vehicle or one of two orally active, MMP-2-preferring inhibitors, ARP-100 or ONO-4817 (10 µM), in SF-188 and A549 cancer cell lines. SF-188 is a human glioblastoma cell line commonly used as a cellular model of pediatric cancers (Rutka et al. 1987), whereas A549 is a model of adult lung carcinoma (Foster et al. 1998). Neither inhibitor, when combined with cisplatin, altered cisplatin’s anticancer activity in SF-188 (Fig. 5A) or A549 (Fig. 5B) cells.

**Figure 5.**
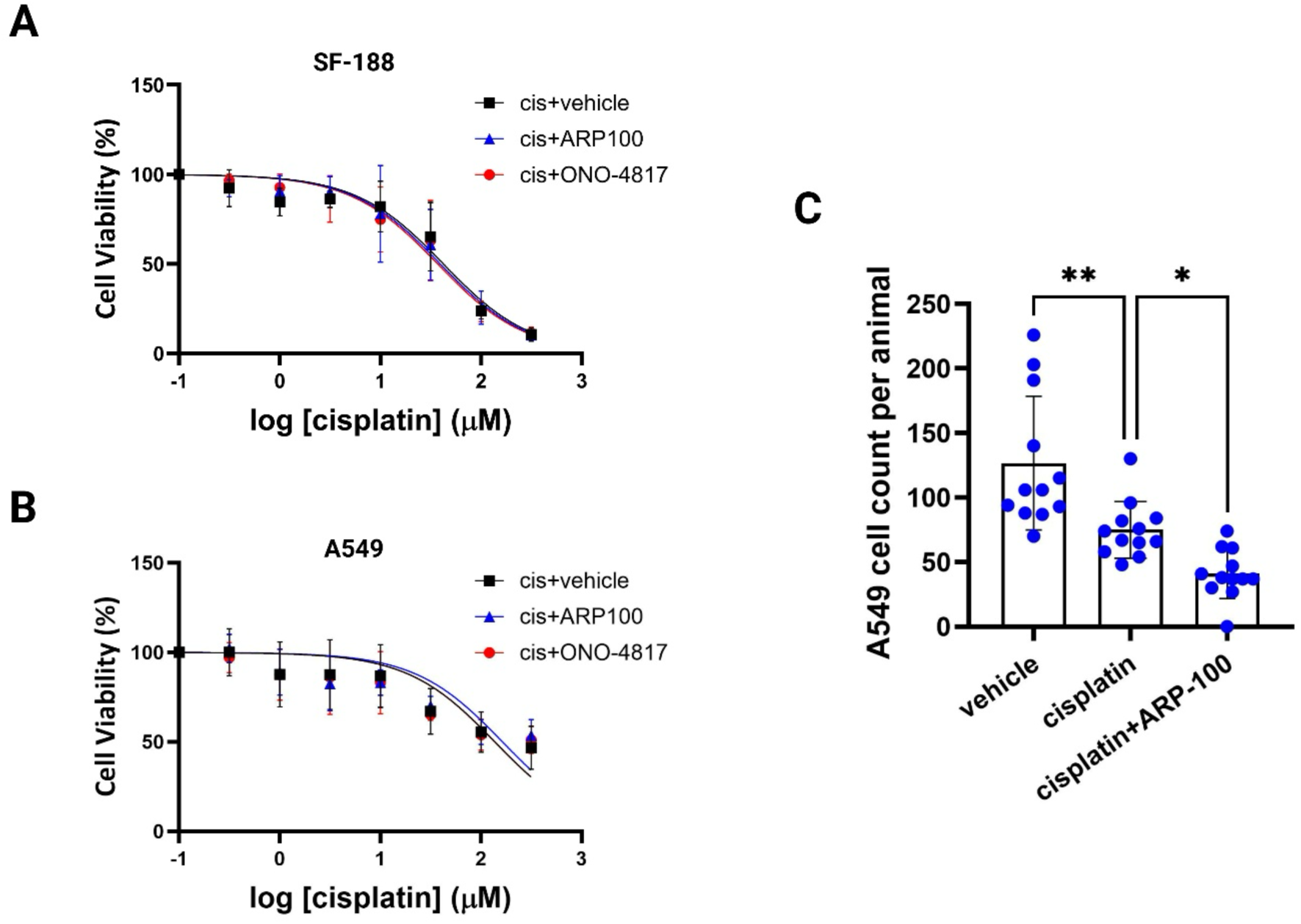
MMP-2-preferring inhibitors do not interfere with the anticancer activity of cisplatin. Cell viability of SF-188 (A) and A549 (B) cells was assessed using the MTT assay after 24 hours of treatment with cisplatin in the presence of vehicle, 10 μM ARP-100, or ONO-4817. The data are represented as the mean ± SD of three independent experiments. C) Fish were injected with A549-mCherry cells at 2 dpf and then bath treated with cisplatin (40μM) with or without ARP-100 (50 μM) or vehicle for 48 h. The number of cancer cells at the end of the treatment was quantified using a confocal microscope. Each dot represents one zebrafish larva, and each treatment group contains 12-13 larvae. Statistical significance was determined using one-way ANOVA followed by Dunnett’s post-hoc test (\**p* < 0.05 and \*\**p* < 0.005).

To determine whether MMP-2 inhibition adversely affects the antitumor efficacy of cisplatin *in vivo*, we compared the therapeutic performance of cisplatin alone with that of cisplatin combined with ARP-100 in a zebrafish cancer xenograft model. To establish a tolerated dose of ARP-100 in zebrafish larvae, we first generated a survival dose-response curve and selected 50 µM as a high dose that was well tolerated (Supplementary Fig. 4). Larval zebrafish were grafted with A549 cells stably expressing mCherry at 2 days post-fertilization. At 1-day post-injection (dpi), larvae were treated with 40 µM cisplatin, cisplatin in combination with ARP-100, or the vehicle added to the bath water. At 3 dpi (48 h post-treatment), larvae were dissociated, and mCherry-positive A549 cells were quantified by confocal imaging and image analysis. As expected, cisplatin significantly reduced the mean tumor burden per animal from an average of 126 (vehicle) to 75 (cisplatin) cells (Fig. 5C). Notably, larvae treated with both cisplatin and ARP-100 exhibited the lowest tumor burden, which remained significantly lower than that of larvae not receiving cisplatin, with an average of 41 cells per larva (Fig. 5C). These data indicate that ARP-100 does not compromise the antitumor efficacy of cisplatin *in vitro* or *in vivo* and may even enhance cisplatin’s therapeutic effect.

### Intracellular MMP-2 mediates a subset of cisplatin-induced proteome changes

To further investigate the proteome changes and putative proteolytic substrates of MMP-2 upon activation by cisplatin, HEI-OC1 cells were pre-treated with 5 μM ARP-100 or vehicle for 2 hours and then co-treated with 20 μM cisplatin and ARP-100 or the vehicle for an additional 6 hours. Cells treated with ARP-100 or vehicle alone were also included as controls. Proteins were extracted from cell lysates and analyzed by Data-Independent Acquisition (DIA) LC-MS/MS. Approximately 7,000 proteins were identified in each lysate that showed a strong overlap, with 6935 proteins being common among the four different groups (Supplementary Fig. 5). Several housekeeping proteins, such as β-2 macroglobulin and β-actin, showed no significant changes in protein abundance between the four groups, as expected.

A comparative analysis of protein abundance levels between cisplatin + vehicle and the vehicle control was performed to explore the effect of cisplatin alone (Fig. 6A). Using an unpaired Student’s t-test (*p*-value <0.05), we measured 236 proteins decreased, and 52 proteins increased in abundance by more than 2-fold. The comparison between cisplatin-treated groups (with or without MMP-2 inhibitor) was also performed to investigate the effect of MMP-2. We found 94 proteins decreased, and 46 proteins increased in abundance by more than 2-fold and *p*-value < 0.05 (Fig. 6B).

**Figure 6.**
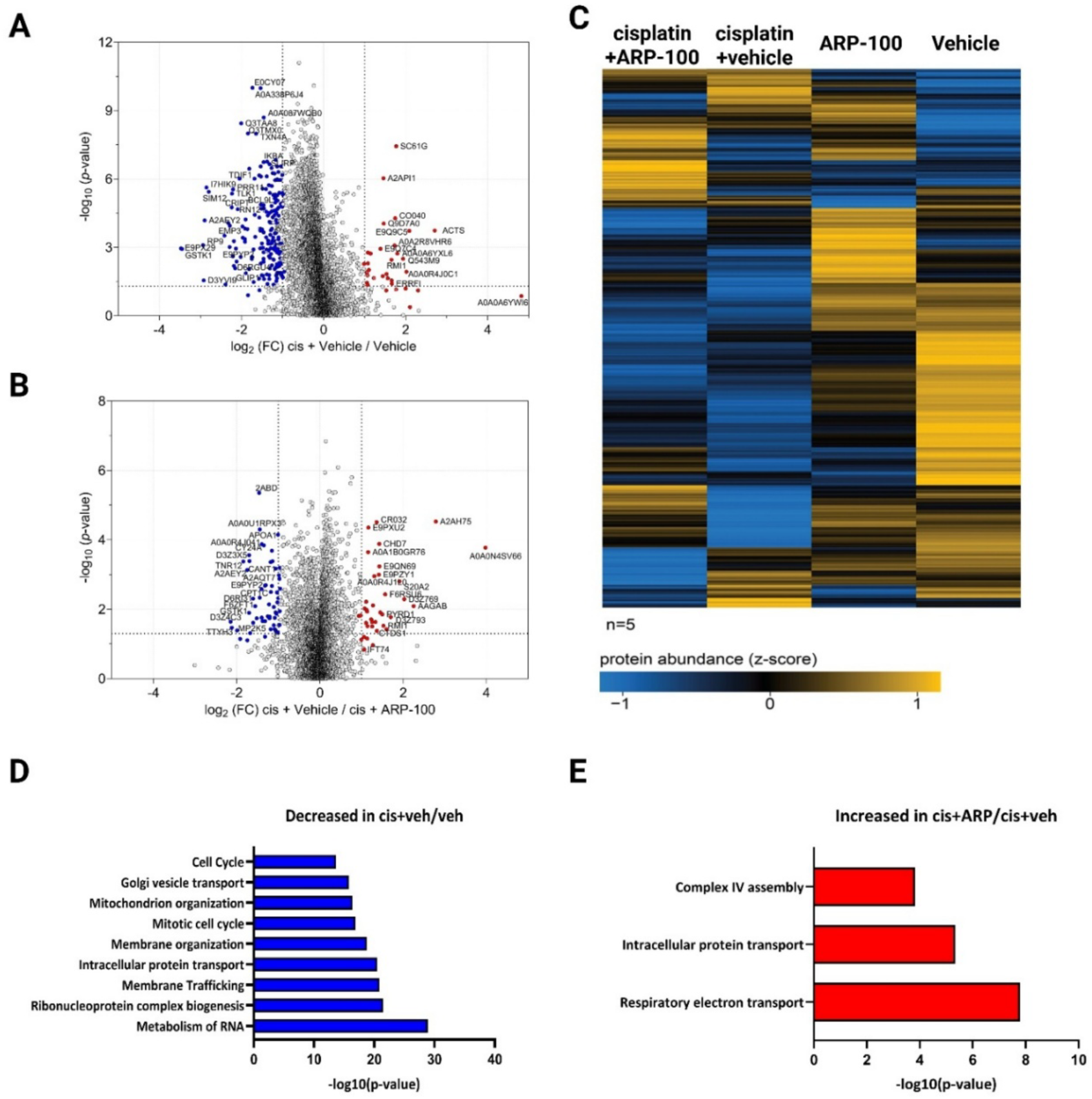
MMP-2 mediates a subset of cisplatin-induced changes in hair cell proteome. HEI-OC1 cells were pre-treated with 5 μM ARP-100 or vehicle for 2 hours and then treated with 20 μM cisplatin for 6 hours. Cell lysates were collected and analyzed by LC-MS/MS. (A, B) Volcano plots comparing protein abundance between the cisplatin + vehicle vs vehicle (A) and cisplatin + vehicle vs cisplatin + ARP-100 groups (B). The Y axis shows negative log_10_ *p-*value, and the x-axis represents proteins with a log_2_ fold change (FC). Red dots represent more abundance, while blue dots show less abundance. (C) Heatmap showing distinct protein abundance patterns across treatment groups. Each column represents a treatment group, and each row corresponds to a protein. Abundance values are scaled using z-scores, with blue indicating lower and orange indicating higher relative abundance. Proteins are grouped by hierarchical clustering using One Minus Pearson correlation. Gene ontology analysis (D, E) shows the enriched biological processes decreased in the cisplatin group (cis+veh/veh, D) or increased in the presence of ARP-100 (cis+ARP/cis+veh, E).

To compare relative changes in protein abundance across experimental groups, a heatmap was generated using z-scores (Fig. 6C). Distinct patterns of protein abundance were observed, but we focused primarily on proteins that were downregulated by cisplatin in the presence of MMP-2 activity (cisplatin + Vehicle) yet upregulated when MMP-2 was inhibited (cisplatin + ARP-100). A gene ontology (GO) analysis of proteins following this pattern was also conducted (Fig. 6D, E). The results showed that proteins involved in cell cycle, mitochondrial organization, membrane organization, membrane trafficking, RNA metabolism, and ribonucleoprotein biogenesis were less abundant after cisplatin exposure. Notably, proteins involved in intracellular trafficking processes were less abundant in the presence of MMP-2 but more abundant upon MMP-2 inhibition, suggesting that they may be direct or indirect substrates of MMP-2.

### MMP-2 degrades RAB proteins after cisplatin treatment

Focusing on proteins involved in intracellular trafficking based on the mass spectrometry and GO analysis, we found that 20 members of the Ras associate binding (RAB) protein family were downregulated by cisplatin, and that MMP-2 inhibition partially restored their abundance (Fig. 7A). RAB proteins are small GTPases that act as molecular switches, regulating vesicular trafficking, directing transport from the plasma membrane to intracellular organelles as well as from intracellular compartments to the secretory pathway (Jordens et al. 2005). This finding is particularly noteworthy because cisplatin enters cells and is actively exported to protect against its toxicity (Safaei, Katano, et al. 2005); disruption of the trafficking system could therefore lead to cisplatin accumulation within cells, exacerbating cellular damage.

**Figure 7.**
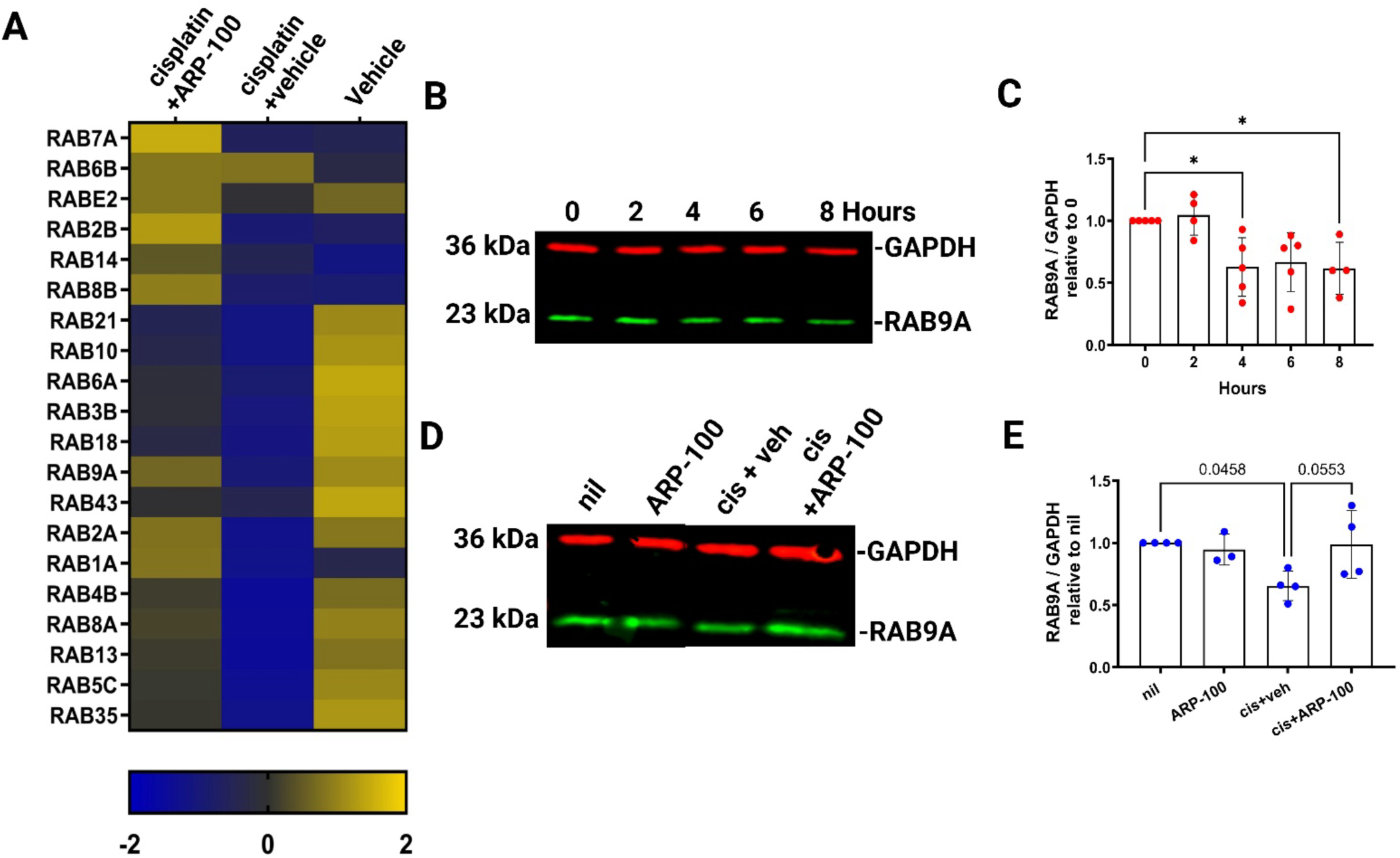
MMP-2 degrades RAB proteins after cisplatin treatment. A) Heatmap showing protein abundance of various RABs in response to cisplatin in the presence or absence of ARP-100. Abundance values are scaled using z-scores, with blue indicating lower and orange indicating higher relative expression. B,C) Immunoblot (B) and quantitation (C) of RAB9A in cytosolic fractions of HEI-OC1 cells treated with 100 μM cisplatin at multiple timepoints. D,E) Immunoblot (D) and quantitation (E) of RAB9A in cytosolic fractions of HEI-OC1 cells pre-treated with or without ARP-100 (10 μM, 2 h) or vehicle, and treated with cisplatin (100 μM, 5 h), or left untreated (nil). The data are represented as the mean ± SD of 3-5 independent experiments. Statistical significance was determined using one-way ANOVA followed by Dunnett’s post-hoc test (\**p* < 0.05).

To validate this finding, we examined RAB9A, a key regulator of protein trafficking from endosomes to the trans-Golgi network and of lysosome biogenesis (Homma et al. 2019). HEI-OC1 cells treated with cisplatin showed a significant ∼50% reduction in RAB9A abundance at 4 and 8 hours compared to untreated controls (Fig. 7B, C), consistent with rapid MMP-2 activation. Pre-treatment with the MMP-2 inhibitor ARP-100 partially restored RAB9A levels (*p* = 0.055; Fig. 7D, E), supporting the mass spectrometry findings.

### RAB9A is proteolyzed by MMP-2

To directly assess whether MMP-2 targets RAB9A, recombinant RAB9A (0.5 μg) was mixed with 100 ng MMP-2 and incubated at 37 °C for various time points. As shown in Fig. 8A, RAB9A protein levels began to decline after 30 minutes of incubation and were completely absent by 90 minutes. In contrast, co-treatment with MMP-2-preferring inhibitors ARP-100 or ONO-4817 restored RAB9A levels (Fig. 8B), supporting RAB9A as a target of MMP-2.

**Figure 8.**
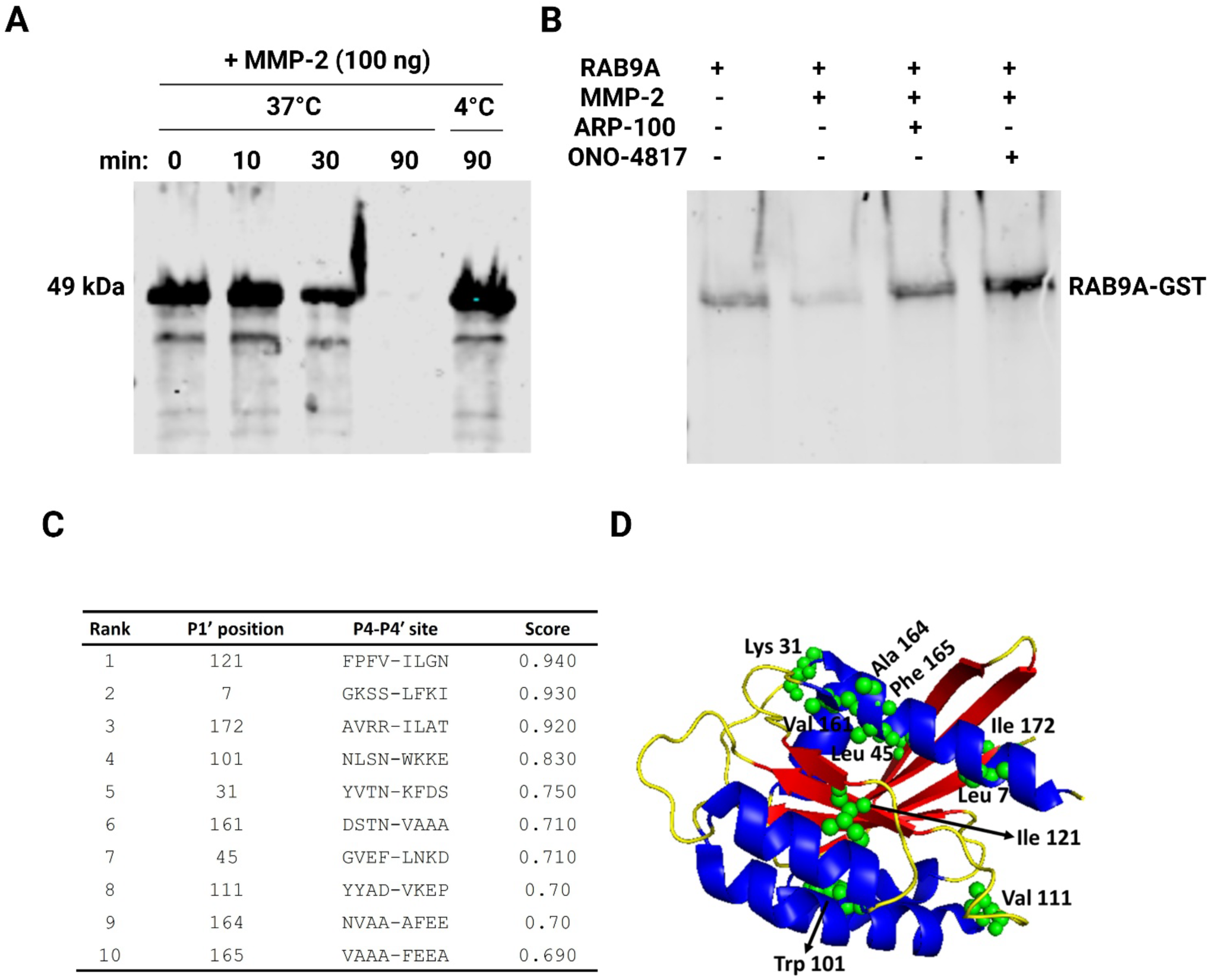
RAB9A is proteolyzed by MMP-2. A, B) Immunoblot of *in vitro* time course proteolysis of recombinant RAB9A at 37 °C with 100 ng MMP-2 (A) or for 30 minutes in the presence or absence of MMP-2-preferring inhibitors (B). C) *In silico* analysis of the top 10 MMP-2 cleavage sites in the structure of mouse RAB9A according to ProsperousPlus. D) Crystal structure of mouse RAB9A showing the predicted cleavage sites (green balls).

Potential MMP-2 cleavage sites on mouse RAB9A were predicted *in silico* using ProsperousPlus. The top 10 predicted sites and their surrounding sequences are shown in Fig. 8C. Four of these sites contain leucine or isoleucine at the P1’ position, consistent with previous studies (Bassiouni et al. 2023; Eckhard et al. 2016). Notably, Lys45 lies within the switch I motif of the GTPase domain, which is critical for conformational changes associated with GTP/GDP binding, while Ile121 resides within the G4 motif responsible for guanine recognition (Wittmann and Rudolph 2004). Mapping these predicted sites onto the 3D structure of mouse RAB9A (Fig. 8D) revealed that some sites, such as Lys31 and Val111, are located in unstructured regions, potentially making them more accessible targets for MMP-2.

## Discussion

In this study, we investigated, for the first time, the impact of MMP-2 and MMP-9 activation on cisplatin-induced ototoxicity, focusing on their roles in cell death, inflammation, and proteomic alterations in cochlear cells. In this *in vitro* work, we tested two orally active compounds that preferentially inhibit MMP-2 and inhibit MMP-9 at higher concentrations (Rossello et al. 2004; Yamada et al. 2000). We found that inhibition of MMPs reduced cisplatin-induced production and secretion of the pro-inflammatory cytokine IL-6, thereby mitigating apoptotic death and improving the viability of auditory hair cells. Furthermore, MMP-2 inhibition could reverse some of the proteome changes induced by cisplatin in auditory cells. Importantly, these inhibitors did not interfere with the anti-cancer activity of cisplatin *in vitro* and even improved it in a zebrafish xenograft model. Currently, sodium thiosulfate is the only FDA-approved drug for preventing CIO; however, its use is limited to patients with non-metastatic localized solid tumors, and concerns remain that early administration may reduce cisplatin’s antitumor efficacy (Freyer et al. 2017). The use of MMP-2-prefering inhibitors to protect against CIO is particularly compelling given the role of MMPs in cancer metastasis (Kleiner and Stetler-Stevenson 1999; Hu et al. 2007). Thus, targeting MMPs could, potentially, provide dual benefits: mitigating ototoxicity while suppressing tumor progression. However, this concept requires further validation *in vivo* using cancer models. While broad-spectrum MMP inhibitors have previously been shown to protect against endotoxin- and noise-induced hearing loss (Jang et al. 2014; Hu et al. 2012; Choi et al. 2012), the specific roles of MMP-2 and MMP-9 in CIO have not yet been addressed.

The role of extracellular MMP isoforms, particularly MMP-9, has been implicated in the disruption of the blood-labyrinth barrier, which leads to the increased cochlear permeability and accumulation of toxins within the cochlea (Jiang et al. 2019). Interestingly, Yang et al. recently reported that co-treatment of cisplatin with antioxidants ferulic acid and ascorbic acid reduced cisplatin-induced toxicity and hair cell death *in vitro* and *in vivo* by downregulating p38 MAPK-mediated MMP-9 expression (Yang et al. 2025). However, the direct impact of MMP-9 inhibition on hair cell viability was not explored. In this study, we demonstrated for the first time that cisplatin exposure results in the secretion and activation of MMP-2 and MMP-9 in auditory hair cells. Full-length secreted MMPs are known to be directly activated by oxidative stress in both extracellular and intracellular environments and are recognized as effectors of oxidative damage. Specifically, MMP-2 is activated by peroxynitrite-induced S-glutathiolation (Viappiani et al. 2009; Wang et al. 2002), while MMP-9 is activated by peroxynitrite and by hypochlorous acid/chloramine-mediated modification of the cysteine switch (Wang et al. 2022). Thus, the activity of these isoforms in our model may result from RONS generated in response to cisplatin or may reflect a secondary activation following cisplatin-induced cell death and the associated oxidative stress observed at 24-48 hours of exposure.

MMP-2 was first shown to exhibit proteolytic intracellular activity in rat hearts during myocardial ischemia-reperfusion injury, where it cleaves troponin I (Wang et al. 2002). MMP-9 has also been detected in intracellular compartments, including the nucleus of neurons and retinal cells, and may contribute to stress responses (Hill et al. 2012; Kowluru et al. 2011), although its functional roles in health and disease are less well-characterized compared to MMP-2. Here, we demonstrated that both MMP-2 and MMP-9 are expressed and rapidly activated in the cytosolic fraction of HEI-OC1 cells following treatment with cisplatin. MMP-2 has been reported in multiple subcellular compartments, including mitochondria, cytoskeleton, nuclei, nucleoli, and the endoplasmic reticulum (Ali et al. 2021; Hughes et al. 2014; Ali et al. 2012; Ren et al. 2019; Kwan et al. 2004). We detected MMP-2 and MMP-9 activity in the mitochondrial fraction, but their activity did not increase in response to cisplatin. In contrast, nuclear levels of MMP-2 and MMP-9 were elevated, suggesting a potential role in intracellular stress responses. Further studies are required to define the precise localization and regulation of MMP-2 and MMP-9 within auditory hair cells.

The intracellular MMP-2 may correspond to the N-terminal truncated (NTT) isoform, which is constitutively active due to the lack of the prodomain and cysteine switch (Lovett et al. 2012). Under basal conditions, the expression of this isoform is very low (or non-existent) but increases under oxidative stress (Lovett et al. 2013). In our study, the *NTT-Mmp-2* expression was upregulated 1 hour after cisplatin exposure. NTT-MMP-2 has been shown in myoblasts to regulate innate immunity by cleaving IκB-α and activating NF-κB (Lovett et al. 2012), which could explain the cisplatin-induced increase in IL-6 secretion and its suppression by MMP-2-prefering inhibitors. NTT-MMP-2 has been shown to specifically induce the transcriptional upregulation of *IFIT-1* (Lovett et al. 2012), a regulator of ribosomal translation. In our study, we observed that cisplatin treatment significantly upregulated *Ifit-1* expression, an effect that was completely reversed by the MMP-2 inhibitor, ARP-100, highlighting the cisplatin-induced activation of NTT-MMP-2. The intracellular activity may also reflect full-length MMP-2 retained in the cytosol and activated by RONS, since up to 50% of nascent MMP-2 fails to be secreted due to insufficient signal peptide efficacy (Ali et al. 2012). Full-length MMP-2 can also modulate innate immunity by cleaving chemokines and cytokines (Song et al. 2013). Although direct evidence for intracellular MMP-9 in auditory hair cells is lacking, studies in neurons suggest that cytosolic MMP-9 may contribute to stress-induced apoptosis and DNA damage (Kimura-Ohba and Yang 2016; Hill et al. 2012). Together, these observations suggest that both intracellular MMP-2 and MMP-9 could contribute to the inflammatory response and apoptosis in cochlear hair cells under cisplatin-induced stress.

Another finding of this study is that MMP-2 downregulates the levels of proteins involved in intracellular trafficking, especially those in the RAB family, upon cisplatin exposure, which might lead to cisplatin retention in auditory cells. We performed an *in silico* prediction study to identify potential cleavage sites of MMP-2 in several RABs, including RAB9A, RAB7, and RAB5. The result showed possible cleavage sites in RABs, highlighting the possible proteolysis of RABs by MMP-2. Although RAB fragments were not detected after incubation with MMP-2, RAB9A protein itself became undetectable, raising the possibility that MMP-2 may modulate the stability of RABs rather than degrading them, or the proteolyzed fragments are not detectable by the antibody. The interplay between RABs and MMPs may be bidirectional. While RABs are involved in the trafficking and secretion of MMPs (Jacob et al. 2013; Stephens et al. 2019), MMP-2 itself appears to modulate RAB protein levels, suggesting a potential regulatory loop. Given that RAB GTPases are essential for intracellular trafficking, organelle homeostasis, and survival (Homma et al. 2019), disruption of their balance could have significant consequences for cell fate. For example, RAB1 and RAB5 are critical for epithelial cell survival, RAB6 deficiency leads to impaired ECM secretion, and RAB7A knockout results in enlarged lysosomes due to lysosomal dysfunction (Homma et al. 2019). RAB9 regulates mannose-6-phosphate receptor transport, and its inhibition causes mis-sorting of lysosomal enzymes (Riederer et al. 1994; Homma et al. 2019), further underlining the importance of RABs in vesicle and lysosomal trafficking.

Several studies have linked RAB proteins to the response to cisplatin through diverse mechanisms, including enhancement of autophagy and mitophagy, inhibition of apoptosis (B. Wang et al. 2023), promotion of cell survival (Ji et al. 2020), and induction of cisplatin resistance via upregulation of the ABCG2 transporter or proliferative pathways (Lian et al. 2017; Zhang et al. 2018). These findings suggest that RABs contribute to cisplatin resistance and cell survival across multiple cancer models. Cisplatin is rapidly sequestered into vesicular compartments and secreted via exosomes (Safaei, Katano, et al. 2005), with resistant cells releasing more cisplatin-containing exosomes than sensitive ones. This process highlights how vesicle trafficking pathways, regulated by RABs, can determine cell survival or death in response to cisplatin (Safaei, Larson, et al. 2005). Although these findings derive mainly from cancer models, RAB proteins are also essential for auditory hair cell integrity and neuronal survival.

In the cochlea, RABs are essential for vestibular hair cell polarity, balance (Chen et al. 2021), and synaptic vesicle trafficking (Heidrych et al. 2008). Therefore, disruption of RAB-mediated trafficking contributes to hearing loss, whereas enhancing vesicular transport can protect against cisplatin toxicity. For example, Waissbluth et al reported that RAB2A and RAB6A were downregulated in the cochlea after cisplatin exposure, whereas treatment with erdosteine, a thiol-based antioxidant, restored their levels and improved cochlear protection (Waissbluth et al. 2017), suggesting that strengthening RAB-mediated vesicular transport and exocytosis represents a compensatory survival mechanism in the CIO condition. Another possible mechanism by which RABs upregulation following MMP-2 inhibition may protect hair cells is through the enhancement of mitophagy. Induction of mitophagy prior to cisplatin treatment has shown promising results in mitigating CIO (Cho et al. 2021), and RABS, including RAB7 and RAB9, are essential for mitophagosome formation and maturation (Homma et al. 2019). Thus, stabilization of RAB by MMP-2 inhibition could promote mitophagy and reduce mitochondrial damage, providing a potential protective mechanism against CIO, although this requires further investigation.

Together, our findings suggest that MMP-2 contributes to CIO by inducing inflammation, cell death, and disruption of endosomal trafficking of cisplatin in auditory hair cells. Thus, MMP-2 inhibition may protect hair cells by reducing inflammation and preserving RAB-mediated trafficking, highlighting a dual mechanism of protection against cisplatin ototoxicity.

## Materials and Methods

### Cell culture and treatments

The murine inner ear cell line HEI-OC1 (a gift from Dr. Federico Kalinec, UCLA) was cultured in Dulbecco’s Modified Eagle Medium (DMEM, Millipore Sigma) supplemented with 10% fetal bovine serum (Millipore Sigma), 100 U/mL penicillin, and 100 µg/mL streptomycin (Thermo Fisher Scientific) at 33 °C in 10% CO₂. Human cancer cell line A549 was purchased from American Type Culture Collection (ATCC) and cultured in DMEM with 10% FBS, 100 U/mL penicillin, and 100 µg/mL streptomycin at 37 °C in 5% CO₂. Human cancer cell line SF-188 was purchased from ATCC and cultured in Eagle’s Minimum Essential Medium with 10% FBS at 37 °C in 5% CO₂. Cells were routinely tested for mycoplasma contamination using a PCR-based detection kit (G238, abm). Cisplatin (Teva) was added to cells 24 h after seeding in fresh medium. Inhibitors, including ARP-100 (5 or 10 µM; 13321, Cayman Chemical), ONO-4817 (5 or 10 µM; a gift from Ono Pharmaceutical, Osaka, Japan), or ethanol (vehicle control), were administered 2 h before cisplatin and maintained during cisplatin exposure. For overexpression studies, the pCMV3-C-FLAG vector encoding human MMP-2 ORF (HG10082-CF, Sino Biological) was transfected into HEI-OC1 using jetPRIME Reagent (Polyplus) according to the manufacturer’s protocol.

### Lentiviral transduction of A549 cells

HEK293T cells were transfected with pMDLg/pRRE (12251, Addgene), pRSV-Rev (12253, Addgene), and pMD2.G (12259, Addgene) lentiviral packaging plasmids as well as pLVpuro-CMV-N-mCherry lentiviral plasmid (123221, Addgene), encoding mCherry fluorescent protein and puromycin resistance cassette. After 72 h, viral supernatants were collected, filtered through a Millex-HV 0.45 µm filter (Millipore Sigma), and concentrated by centrifugation. A549 cells were transduced with the prepared virus in DMEM supplemented with 10 µg/mL polybrene. After 72 h, the medium was replaced with selection medium containing puromycin (1–1.5 µg/mL), and mCherry-positive cells were used for subsequent experiments.

### Gelatin zymography

Cells (2 × 10⁵) were seeded in 6-well plates and cultured overnight. The following day, cells were exposed to either increasing concentrations of cisplatin for 24 or 48 h, or to 20 μM cisplatin for 6 h, in serum-free medium. Conditioned media were collected, mixed with non-reducing loading buffer, and separated on 10% polyacrylamide gels copolymerized with 2 mg/mL porcine gelatin type A (G8150, Sigma-Aldrich). After electrophoresis, gels were washed three times with 2.5% Triton X-100 at room temperature for 20 min each, then incubated in the zymography activity buffer (50 mM Tris–HCl, 150 mM NaCl, 5 mM CaCl₂·2H₂O, 0.05% NaN₃, pH 7.6) at 37 °C for 20 h. Gels were subsequently stained with 0.05% Coomassie Brilliant Blue G-250 (Bio-Rad) for 4 h and destained overnight in 4% methanol and 8% acetic acid. For intracellular MMP activity, cytosolic and mitochondrial fractions were prepared using a mitochondrial isolation kit (89874, Thermo Fisher Scientific), following the manufacturer’s protocol. 10 μg of protein were resolved on gelatin-containing gels as described above. Conditioned serum-free medium from HT1080 cells was used as a standard for MMP-2, while medium from cells treated with 100 ng/mL phorbol myristate acetate served as the MMP-9 standard as previously performed (Bassiouni et al. 2023).

### Real-time quantitative PCR

Cells (1 × 10⁵) were seeded in 12-well plates and treated with 20 μM cisplatin for 1, 2, 4, or 24 h, or left untreated. Total RNA was extracted using the Aurum Total RNA kit (7326820, Bio-Rad) following the manufacturer’s protocol. cDNA was synthesized from 500 ng of total RNA using the iScript cDNA Synthesis kit (1708890, Bio-Rad). Gene expression of *B2M*, full-length *Mmp-2*, N-terminal truncated *Mmp-2*, and *Mmp-9* was assessed with the primers listed in Supplementary Table 1 using SYBR Green PCR Master Mix (A25742, Applied Biosystems). Expression of *B2M* (Mm00437762_m1), *Il-6* (Mm00446190_m1), and *Ifit-1* (Mm00515153_m1) was measured using validated Applied Biosystems TaqMan assays in combination with the SsoAdvanced Universal Probes Supermix (1725281, Bio-Rad). RT-qPCR was performed on an Applied Biosystems real-time PCR system, and relative expression levels were calculated using the 2^−ΔΔCq^ method.

### Immunofluorescent confocal microscopy

Cells (3 × 10⁴) were plated on 12 mm German glass coverslips (Electron Microscopy Sciences) in 24-well plates and cultured in DMEM overnight. The following day, cells were treated with 20 μM cisplatin for 6 h or left untreated. Cells were fixed with 4% paraformaldehyde (Thermo Fisher Scientific) for 10 min at room temperature, washed three times with PBS (5 min each), and permeabilized with 0.1% Triton X-100 in PBS for 15 min. After an additional PBS wash, cells were blocked in 5% FBS in PBS for 30 min at room temperature. Primary antibody incubation was performed overnight at 4 °C in PBS containing 0.5% FBS and 0.1% Tween-20 with either rabbit monoclonal anti-MMP-2 (1:200, ab92536, Abcam) or mouse monoclonal anti-MMP-9 (1:50, sc-393859, Santa Cruz Biotechnology). The next day, cells were washed three times with PBS and incubated for 1 h at room temperature with secondary antibodies prepared in PBS containing 0.5% FBS and 0.1% Tween-20, and Alexa Fluor 647 goat anti-rabbit (1:500, 111605003, Jackson ImmunoResearch) or Alexa Fluor 488 goat anti-mouse (1:500, 4408S, Cell Signaling). The cells were then washed twice with PBS and once in PBS containing 0.5μg/mL DAPI and two more times with PBS at room temperature before mounting using 1 drop of Prolong Glass Antifade Mountant (Invitrogen). Slides were allowed to dry overnight at room temperature, away from light. Imaging was performed using a spinning disk confocal microscope (Quorum Technologies) at the University of Alberta Cell Imaging Core, equipped with the appropriate emission filters for DAPI, GFP, and Cy5, and a 60×/1.42 NA oil-immersion objective. Images were analyzed using Fiji software.

### Cell viability assay

HEI-OC1 (5 × 10³) or SF-188 (10× 10³) cells were seeded in 96-well plates and treated for 24 h with increasing concentrations of cisplatin (0.316–316 μM) in the presence or absence of inhibitors or vehicle. 1 mg/mL of the MTT reagent (M6494, Invitrogen) was then added, and plates were incubated for 4 h at 33 °C with 10% CO₂ in the dark. After incubation, the media was removed and replaced with dimethyl sulfoxide. The plates were shaken at room temperature until the formazan crystals were fully dissolved. Absorbance was measured at 590 nm using a SpectraMax i3x plate reader (Molecular Devices). Cell viability was calculated relative to untreated controls, defined as 100% viable, using the formula:

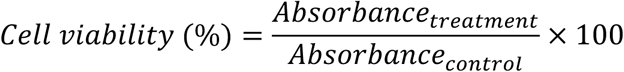

### Flow cytometry

Cells (1 × 10⁵) were seeded in 12-well plates and treated for 24 h with 20 μM cisplatin in the presence or absence of 5 μM ONO-4817 or vehicle control. Cells were harvested by trypsinization, washed twice with Annexin V binding buffer (BMS500FI, Invitrogen), and stained with Annexin V–FITC (1:20) for 15 min at room temperature. After two additional washes with binding buffer, cells were mixed with propidium iodide (PI; 1:40). For each condition, at least 10,000 events were collected on an Attune NxT flow cytometer (Thermo Fisher Scientific) at the University of Alberta Flow Cytometry Core. Data were analyzed using FlowJo software (BD Biosciences).

### Enzyme-linked immunosorbent assay (ELISA)

Cells (5 × 10⁴) were seeded in 24-well plates and treated for 24 h with 20 μM cisplatin in the presence or absence of 5 μM ONO-4817, ARP-100, or vehicle. In parallel experiments, cells were transfected with an MMP-2-expressing vector, an empty vector, or siRNAs 24 h prior to cisplatin treatment. Following treatment, culture supernatants were collected (leaving adherent cells for MTT assay). IL-6 levels were quantified using a mouse IL-6 ELISA kit (88706488, Invitrogen) according to the manufacturer’s instructions. Absorbance was measured at 450 and 570 nm on a SpectraMax i3x plate reader (Molecular Devices). IL-6 secretion was normalized to the number of viable cells, determined by the MTT assay, to account for differences in cytotoxicity across treatment groups.

### siRNA gene knockdown

Cells (5 × 10⁴) were seeded in 24-well plates and cultured overnight to reach ∼50–60% confluency. The next day, cells were transfected with either non-targeting control siRNA or Mmp-9 siRNA (a pool of three siRNAs targeting different regions; mm.Ri.Mmp9.13.1-3) from the TriFECTa RNAi Kit (Integrated DNA Technologies) using jetPRIME transfection reagent (Polyplus). After 4 h, the medium was replaced with fresh medium, and cells were incubated for an additional 24 h before treatment with 20 μM cisplatin for 24 h. Media were collected for IL-6 ELISA, and adherent cells were used for the MTT assay for normalization, as described above.

### Immunoblotting

Cells were lysed in NP-40 cell lysis buffer supplemented with protease inhibitor tablets (A32963, Thermo Fisher Scientific). Equal amounts of protein (20 μg) were separated on 12% SDS–polyacrylamide gels, wet transferred to nitrocellulose membranes, and blocked with Intercept (TBS) Blocking Buffer (92760001, LI-COR) for 1 h at room temperature. Membranes were probed overnight at 4 °C with the following primary antibodies: mouse monoclonal anti-GAPDH (1:2500, MA515738, Invitrogen), rabbit polyclonal anti-RAB9A (1:1000, 114201AP, Proteintech), and rabbit monoclonal anti-MMP-2 (1:1000, ab92536, abcam). After washing with TBST, membranes were incubated with IRDye 800RD goat anti-rabbit IgG (92568071, LI-COR) or IRDye 680RD goat anti-mouse IgG (92668020, LI-COR) secondary antibodies for 1 h at room temperature. Blots were visualized using the Odyssey Infrared Imaging System (LI-COR), and band intensities were quantified with ImageJ software. Protein levels were normalized to GAPDH as a loading control.

### Liquid chromatography tandem mass spectrometry (LC–MS/MS)

Cells (10⁶ per experimental group) were lysed in RIPA buffer supplemented with 5% SDS. Proteins from four biological replicates (50 µg each) were acidified with 1.2% (v/v) phosphoric acid at room temperature and subsequently captured using Suspension-Trapping columns (S-Trap, ProtiFi). Columns were washed four times with 90% methanol in 100 mM tetraethylammonium bicarbonate to remove residual contaminants. The retained proteins were reduced, alkylated, and digested overnight at 37 °C with mass-spectrometry grade trypsin (Promega) at a 1:10 enzyme-to-protein ratio. Resulting peptides were sequentially eluted with 0.2% formic acid and 50% acetonitrile, then vacuum dried. Samples were reconstituted in 0.1% trifluoroacetic acid, desalted using C18 ZipTips (Millipore Sigma), dried again, and finally resuspended in 0.1% formic acid for LC-MS/MS analysis.

Mass spectrometry was carried out using reverse phase LC (Thermo Scientific EASY-nLC 1200) interfaced to an Orbitrap Fusion Lumos Tribrid (Thermo Fisher Scientific) mass spectrometer. A 120-minute linear gradient ranging from 3.85% to 36.8% acetonitrile in 0.1% formic acid was applied for eluting the peptides. Data were acquired in data-independent acquisition (DIA) mode with MS1 recorded in the Orbitrap at 120,000 resolution (FWHM) across an m/z range of 350-2,000. MS2 scans were collected at 30,000 resolution over 350-1,400 m/z using an isolation width of 38.5 m/z. Higher-energy collision dissociation (HCD) was employed with a positive ion voltage of 1,650 V. Dynamic exclusion was enabled, with ions excluded for 45 seconds following a single detection event.

Raw spectral data were processed in Spectronaut software (version 18) using the mouse reference proteome obtained from UniProt (https://www.uniprot.org/). To control for false discoveries, peptide identifications were compared against a decoy database, with the false discovery rate threshold set at 1%. For visualization, protein abundance values were z-score normalized across samples, and heatmaps were generated in R (v4.2.1; R Core Team, 2018) using the *superheat* package. Statistical significance for comparisons was assessed by two-way ANOVA followed by Tukey’s honest significant difference post hoc test. Gene ontology analyses were performed using https://metascape.org against the mouse proteome on proteins with significant changes in abundance (*p* < 0.05).

### *In vitro* proteolysis of RAB9A by MMP-2

0.5 μg of recombinant human RAB9A-GST fusion protein (Ag1969, Proteintech) was incubated with 100 ng of recombinant MMP-2 Fc-fusion protein containing the catalytic domain of human MMP-2 (Hartley et al. 2024) for 10, 30, and 90 min in the presence or absence of ARP-100 or ONO-4817 (30 μM) at 37°C in activation buffer (50 mM Tris–HCl, 150 mM NaCl, 5 mM CaCl2, pH 7.6). The reaction was stopped by mixing with the loading buffer. The samples were then separated on a 12% SDS-polyacrylamide gel, transferred to nitrocellulose membrane, and probed with an anti-RAB9A antibody as previously described. An additional mixture of MMP-2 and RAB9A was kept at 4°C to rule out any non-specific reaction.

### *In silico* analysis of RAB9A cleavage sites by MMP-2

The online cleavage prediction server ProsperousPlus was used to predict the top ten potential cleavage sites of MMP-2 on the amino acid backbone of RAB9A. This server suggests the *in silico* prediction of cleavage sites of a variety of enzymes. The mouse RAB9A FASTA sequence (UniProt ID-Q9R0M6) was used as the query, while MMP-2 was selected as the enzyme of interest.

## Crystal structure analysis of MMP-2 cleavage sites in RAB9A

The amino acid sequence of mouse RAB9A was obtained from UniProt, and its X-ray crystal structure was retrieved from the Protein Data Bank (PDB). Mouse RAB9A consists of 201 amino acids; however, the X-ray structure spans residues 2–175. Because the top ten predicted MMP-2 cleavage sites fell within this range, the X-ray structure was used for analysis rather than the AlphaFold model. The top ten predicted cleavage sites were mapped and visualized on the RAB9A structure using PyMOL.

### Animal ethics and zebrafish husbandry

Husbandry and breeding of zebrafish were approved under the protocol AUP00000077 by the Animal Care and Use Committee, BioSciences at the University of Alberta, under the auspices of the Canadian Council on Animal Care. Adult zebrafish were maintained according to standard procedures in a 14:10 light cycle in brackish water (1250 ± 50 µS) at 28.5°C and fed twice daily with either brine shrimp or juvenile trout chow. Zebrafish larvae were maintained in embryo media at 28.5 °C in a 14:10 light: dark incubator until the start of experimental procedure (*Westerfield 2000*).

### Cancer Xenograft into Zebrafish Larvae and Drug Treatments

A549-mCherry cells were maintained under conditions as described above. On the day of injection, cells were loaded into glass capillary tubes that had been pulled to use for injection. Larvae at 2 days post-fertilization (dpf) were anaesthetized with 4% tricaine (MS-222, Sigma-Aldrich) and placed in a dish coated with 1.5% agarose, molded to form channels for ease of larval manipulation. Injection volumes were calibrated to deliver 75-100 cells per injection, and A549-mCherry cells were injected into the yolk sac. Larvae were subsequently removed from anaesthesia, recovered for 30-60 minutes at ambient temperature, and incubated at 33 °C, midway between cell and zebrafish optimal temperatures.

At 1-day post-injection (dpi), larvae were screened under an epifluorescence microscope for the presence of mCherry-positive cells, then divided randomly into three drug bath treatment groups: vehicle control (0.1% DMSO, 0.002% ethanol diluted in embryo media), 40 µM cisplatin (in vehicle solution), and cisplatin (40µM) + ARP-100 (50 µM in vehicle solution). DMSO was added to facilitate drug uptake into zebrafish larval tissues (Yan et al. 2025), and ethanol was included as it is the solvent of ARP-100. Larvae were incubated at 33 °C in darkness to prevent cisplatin degradation, and drug baths were changed daily.

### Cell Dissociation and Imaging

At 3 dpi, larvae were euthanized with an overdose of 4% tricaine and underwent cell dissociation, modified from previous protocols (Azzam et al. 2025; Jones et al. 2023). Sample groups of 5-20 larvae were incubated in 1mg/mL of collagenase P (11213857001, Millipore Sigma) at 37 °C for 30 min, with mechanical dissociation at 0, 10, 20, and 30 min via pipetting. Next, 10% normal goat serum/PBS was added to each sample to stop collagenase activity, and samples were gently vortexed and pelleted by centrifugation at 300 *g* for 15 min at 4 °C. The supernatant was removed, and samples were resuspended in 0.04% NGS/PBS. Samples were stored on ice and imaged within 2 hours of dissociation. Dissociated samples were imaged in 10 µL increments on a Leica Stellaris confocal microscope (Leica Microsystems) for detection of mCherry fluorescence. Tiled images were obtained for each 10 µL volume. Negative (uninjected) controls were prepared and imaged in parallel.

Image analysis was performed in Fiji. Tiles were stitched into their composite images using the Grid/Collection Stitch, and thresholding was done in the mCherry channel to reduce background signal. mCherry-positive cells were counted using the Analyze Particles function, with parameters for particle size and circularity. These parameters were established using images of positive controls (A549 mCherry cells present) and negative controls (uninjected, dissociated zebrafish) to eliminate false positives from autofluorescent zebrafish cells.

### Statistics

Data are shown as the mean and standard deviation (SD) of N independent experiments. Each independent experiment corresponds to a distinct passage number derived from the same original cell batch. One-way or two-way ANOVA followed by Dunnett’s post-hoc test was used for multiple comparisons (GraphPad Prism version 10, La Jolla, CA, USA), while p < 0.05 was considered statistically significant.

## Acknowledgements

This work was supported by operating grants to APB from the Canadian Institutes of Health Research (Grant PJT-178327) and the Kids with Cancer Society through the Cancer Research Institute of Northern Alberta, as well as to RS from the Canadian Institutes of Health Research (Grant FDN143299). Z.Z. was supported by studentships from the University of Alberta Faculty of Medicine & Dentistry, the Li Ka Shing Institute of Virology, and the Women and Children’s Health Research Institute. MGD is supported by a Clinical Fellowship from ALS Canada and Brain Canada. Funding to MGD and WTA was from the Alberta Cancer Foundation Palliative Care Pilot Research Grant administered by the Cancer Research Institute of Northern Alberta. APB holds a Canada Research Chair in Pattern Recognition Receptor Pathophysiology and this research was undertaken, in part, thanks to funding from the Canada Research Chairs Program (101342).

Some experiments were performed at the University of Alberta Faculty of Medicine & Dentistry Flow Cytometry Facility (RRID: SCR_019195), which receives financial support from the Faculty of Medicine & Dentistry and Canada Foundation for Innovation (CFI) awards to contributing investigators. Immunofluorescence experiments were conducted at the University of Alberta Faculty of Medicine & Dentistry Cell Imaging Core (RRID: SCR_019200), which is supported by the Faculty of Medicine & Dentistry, the University Hospital Foundation, Striving for Pandemic Preparedness – The Alberta Research Consortium, and CFI awards to contributing investigators. We also thank the Advanced Microscopy Unit and Dr. Kacie Norton for use of imaging resources and support, Dr. Jason Berman and Nadine Azzam for training and support in xenografting, Melissa Kinley for technical support, and the Aquatics Facility Staff for their diligent care of the animals.

## Author contributions

Conceptualization: ZZ, BH, WB, MGD, SL, MJS, WTA, OJ, RS, APB; Investigation: ZZ, BH, WB, MGD, SL; Formal Analysis: ZZ, BH, WB, MGD, SL, APB; Funding acquisition and Supervision: WTA, RS, APB; Writing – original draft: ZZ, BH, MGD; Writing – review & editing: all authors.

