## Supplemental Figs and Table for "Matrix metalloproteinases proteolyze RAB proteins and contribute to cisplatin-induced ototoxicity"

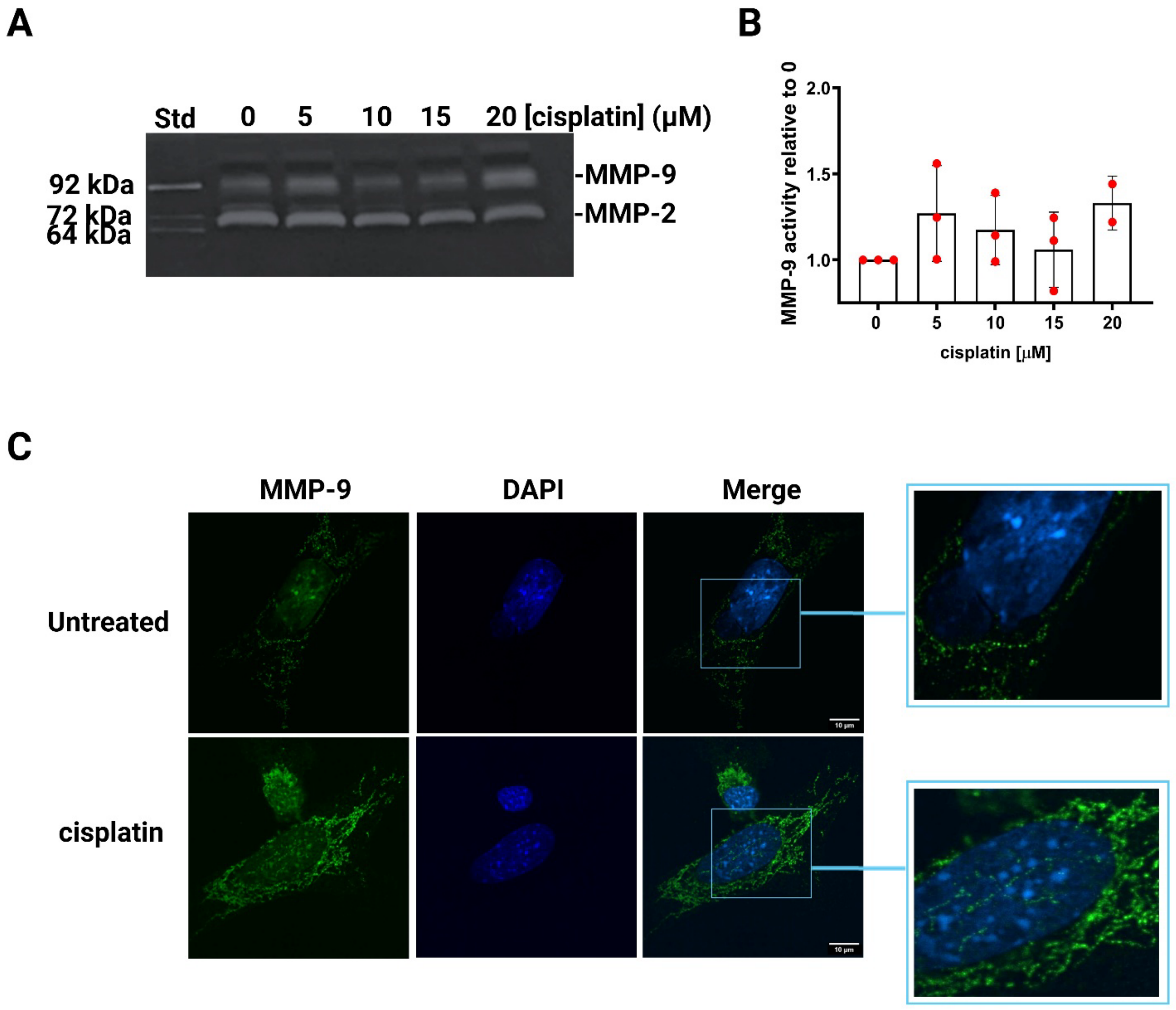


**Supplementary Figure 1. MMP-9 activity and presence in HEI-OC1 cells.** Gelatin zymography (A) and corresponding quantitative analysis (B) of extracellular (secreted) MMP-9 activity in HEI-OC1 cells following 24 h of exposure to 20 µM cisplatin. Std derives from conditioned medium from HT-1080 cells. C) HEI-OC1 cells were treated with 20 µM cisplatin for 6 h or left untreated. Immunofluorescence staining was performed using an antibody specific to MMP-9, followed by detection with a secondary antibody conjugated to Alexa Fluor 455 (green). The zoomed-in boxes indicate nuclei with an increased green signal.

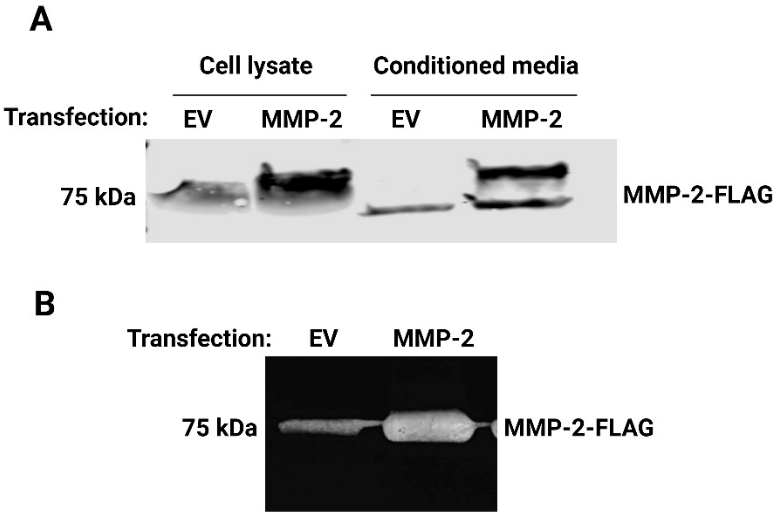


**Supplementary Figure 2. Human MMP-2-FLAG tagged expressing plasmid is successfully expressed and activated in HEI-OC1 cells.** Cells were transfected with a pCMV3 plasmid encoding *Mmp-2* or the empty vector (EV) for 24 h. A) The immunoblot of MMP-2-FLAG (blotted with the anti-MMP-2 antibody) in the cell lysates and conditioned media, and B) Gelatin zymography of MMP-2 activity in the conditioned media of transfected HEI-OC1 cells.

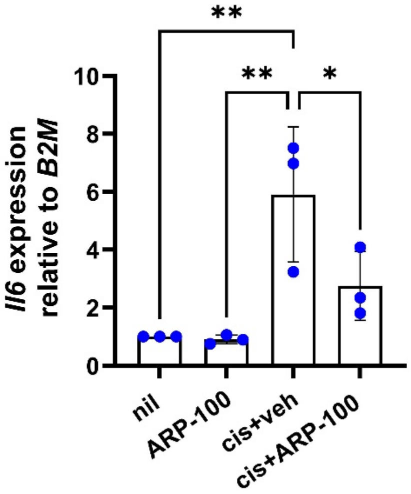


**Supplementary Figure 3. MMP-2-preferring inhibitor attenuates cisplatin-induced *Il-6* expression in HEI-OC1 cells.** mRNA expression of *Il-6* was measured following a 24h exposure to 20 μM cisplatin in the presence or absence of 5 μM ARP-100, compared to the untreated control (nil), and normalized to the expression of the housekeeping gene, *B2M.* The data are represented as the mean ± SD of three independent experiments. Statistical significance was determined using one-way ANOVA followed by Dunnett's post-hoc test (*$p < 0.05$ and **$p < 0.01$).

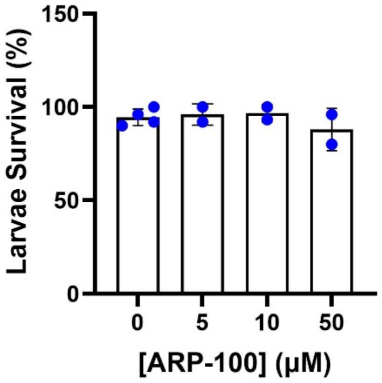


**Supplemental Fig. 4. Zebrafish larvae survival in response to ARP-100.** Each replicate contained more than 15 larvae from 3 days post-fertilization, incubated at 33 °C. Larvae were exposed to the indicated concentrations of ARP-100, and their survival was monitored after 48 hours of treatment.

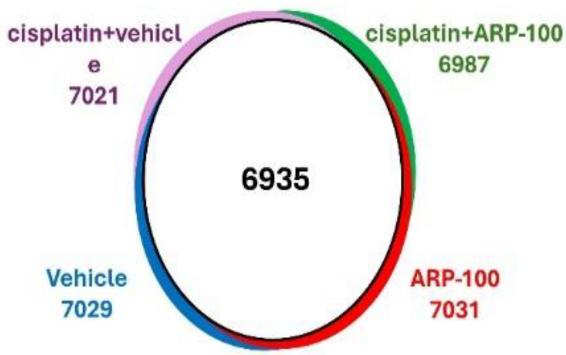


**Supplementary Figure 5.** The proteome data show a strong overlap between the proteins quantified in each group. 6935 proteins were identified at least in one replicate of each group. Less than 8 proteins were found in only one of the treatment groups, as indicated by the small colored area for each.

**Supplementary Table 1.** Primer sequences used for qPCR.

| Gene | Forward Sequence (5' to 3') | Reverse Sequence (5' to 3') | Product size |
|---|---|---|---|
| *B2M* | TGGTCTTTCTGGTGCTTGTCTC | CCCGTTCTTCAGCATTTGGATTTC | 170 bp |
| *Mmp-2* | GACCTCTGCGGGTTCTCTGC | TTGCAACTCTCCTTGGGGCAGC | 163 bp |
| *NTT-Mmp2* | GTGAATCACCCCACTGGTGGGTG | TTGCAACTCTCCTTGGGGCAGC | 229 bp |
| *Mmp-9* | CCTGGCTCTCCTGGCTTTC | ATAGCGGTACAAGTATGCCTCTG | 136 bp |
